# Synthesis, characterization and evaluation of cell culture study of mercury complex derived from benzene-1,3,5-tricarboxylic acid

**DOI:** 10.1101/2023.05.25.542356

**Authors:** Javed Hussain Shah, Shahzad Sharif, Rashid Rehman, Anum Arooj

**Author notes:** **Corresponding Author:** Javed Hussain Shah, Contact No., Postal Address: Materials Chemistry Laboratory, Department of Chemistry, Government College University, Katchery Road Lahore, 54000 Pakistan.

## Abstract

Mercury complexes have diverse effects on the human body and cells that depend upon the biochemical form of mercury-complexes and the nature of exposure. In the present work, we have investigated the impacts of mercury-complex derived from benzene-1,3,5-tricarboxylic acid on cell culture and DNA damage. This is novel mercury-complex having cell culture study. The mercury complex has been synthesized and characterized by CHNS analyzer, FTIR, X-Ray Diffraction (XRD) and DNA damage. Surface morphology of prepared mercury-complex was studied by microscopy imaging/Atomic Force Microscopy (AFM). The main goal of this contribution is to address the damaging effects of mercury-complex in cell cultures through fluorescence imaging and identifying cell Live/Death quantitative results. These live/death color intensities are red or green in presence to the mercury-complex. For this purpose, we measured the concentration dependence of mercury-complex on the rate of death in cells which may be useful for the cell culture and DNA study. The mercury-complex derived from benzene-1,3,5-tricarboxylic acid has the ability to break the polynucleotide structure of DNA to mono nucleotides resulting irreparable DNA damage. The experimental results of fluorescence microscopy and live/dead cell assay on cell viability reflected the potent cellular toxicity of mercury-complex causes cell culture study. Generally, the damaging effects of mercury-complex may be used for medical treatment of different diseases specially cancer.

## Introduction

When confronted with the term “heavy metals,” the general public with any scientific background, and even many chemists, immediately associate it with toxicity and dangerous materials. In fact, few people are aware that an increasing number of compounds containing a heavy-metal ion in a critical role are extremely useful, and several of them are used as drugs to cure or diagnose disease (1). Some heavy metals are required in trace amounts to improve enzyme function and other intracellular molecules (2). Many heavy metals (micronutrients: Cu, Zn, Fe, Mn, Mo, Ni, and Co) are essential for plants and animals when present in low concentrations in the growing medium; those that are becoming noxious only when a concentration limit is crossed (3). Similarly, arsenic, which can be poisonous, and its chronic exposure from industrial or natural sources can cause serious toxicity, at the same time arsenic has been used therapeutically for more than 2,400 years. In the 1930s, arsenic was reported to be effective in chronic myelogenous leukemia (4). Classical, well-known examples include silver compounds (used to protect the skin after burning wounds), radioactive technetium compounds (used as disease diagnostics), bismuth salts (used in the treatment of diarrhea and stomach ulcers), gold complexes (used to treat arthritis), copper salts (also used to treat arthritis), and platinum compounds (which are used as efficient anti-tumor drugs). In fact, depending on the dose, all such metal compounds are poisons; however, some of the most toxic metals are essential for life, either as a trace element or as a component of a drug (1).

For thousands of years, metals and metal compounds have been used in medicine. The ten most active metals in terms of anti-cancer activity are arsenic, antimony, bismuth, gold, vanadium, iron, rhodium, titanium, gallium, and platinum. Arsenic, the first metal examined, has the unusual property of having both anti-cancer and oncogenic properties. Some antimony derivatives, such as Sb_2_O_3_, salt (tartrate), and organic compounds, exhibit intriguing properties. Bismuth directly affects helicobacter pylori and gastric lymphoma; bismuth complexes of 6-mercaptopurine show promising results. Although gold(I) and gold(III) compounds have anti-tumor properties, their toxicity remains high. Gold derivatives are still being researched for their potential use. Several vanadium derivatives exhibit anti-proliferative activity, but their toxicity must be overcome. Several pieces of evidence suggest that iron deficiency could be an excellent therapeutic approach; additionally, it is synergistic with standard anti-cancer drugs. Rhodium belongs to the same group as platinum and exhibits similar activity, but with the same nephrotoxicity. A number of rhodium compounds have begun phase I clinical trials. Titanium derivatives, in contrast to platinum complexes, showed no evidence of nephrotoxicity or myelotoxicity; titanocene dichloride is currently in clinical trials. Gallium’s anti-proliferative effect could be attributed to its competition with the iron atom; additionally, a derivative appears to reverse multidrug resistance. Platinum, the last metal examined, has produced some of the most effective anti-cancer drugs. In the clinic today, four derivatives are used; their mechanisms of action and resistance are described (5).

For centuries, Mercury was knows as one of the most hazardous element (6,7) that leads to the malfunction of cells and consequently causes much damage to the brain, kidney, central nervous system (8,9) and to the environment (10,11). Despite of its severe toxic nature, mercury has been used in different technologies to support human. Such as, three isotopes Hg-198, Hg-200 and Hg-202 are used to find out the entry route of mercury into the ecosystem that affects the fishes. Hg-303 is used for the calibration of gamma radiations while Hg-202 used to produce Hg-303 which is radioactive isotope of mercury (12–14). Previously, Shenker et.al 1993 examined both the immunotoxic and cytotoxic properties of mercuric compounds on human B-cells and their results indicate that low doses of mercury are capable of inhibiting B-cell activation. Unlike T-cells, these inhibitory effects are independent of monocytes (15). Based on above facts, the current study has focused on use of mercury in medical science to open the new avenues to use this compound as therapeutic agent for different diseases.

Currently, all forms of mercury were considered to be immunosuppressive, given their potent cytotoxicity in culture and high-dose effects in animals. Both inorganic mercury compounds HgCl_2_ and methyl mercury (MeHg) prevented mitogen-induced proliferation in B cells and immunoglobulin (IgG and IgM) production *in vitro*, with MeHg being 10 times more potent than HgCl_2_ (15,16). Mercury forms a very stable complex with thio amino acids such as cystein and methionine because of their great affinity towards sulphur (10,11). This metal has a high affinity for enzymes and proteins with thiol groups, which are involved in normal cellular defense mechanisms (17).

Mercury is a strong phytotoxic as well as genotoxic metal (18,19). In terms of genotoxicity, mercuric ions tend to form covalent bonds with DNA. It is known that both organic and metallic mercury have the ability to penetrate into the central nervous system (CNS) (20). There are seven isotopes of mercury which are very stable, and are mainly used in the deposition and emission in both aquatic and terrestrial environment (21). The most naturally occurring mercury isotope is Hg-202, its natural abundance is 29.86% and the least naturally occurring is Hg-196 which has natural abundance 0.15% (12–14). Inorganic mercury makes bond with elements like chlorine, fluorine or any other. These are mostly in white crystalline form and called mercury salts, there is only one exception which is cinnabar (HgS), it is red in color but when it is exposed to light it becomes black in color. Some mercury compounds are violently poisonous such as mercury chloride (HgCl_2_). The most common example of organic mercury is methyl mercury (MeHg), which is present abundantly in environment, when free mercury enters into body, some microorganisms over there convert mercury into methyl mercury (MeHg) (22,23).

HgCl_2_-induced DNA damage has many similarities to those caused by X-rays. However, the single strand breaks induced by HgCl_2_ are not readily repaired, in contrast to those induced by X-rays. HgCl_2_ has been shown to induce frank single strand breaks, not alkali-labile sites (24). DNA repair was assessed by the disappearance of DNA lesions as evaluated by alkaline elution studies and also by CsCl_2_ density gradient analysis. Similar to X-rays, HgCl_2_ has also been shown to cause the formation of superoxide radicals and the depletion of reduced glutathione in intact cells. The binding of mercury to DNA was shown to be very tight since it resisted extraction with high salt and chelating agents (25,26).

Previous researches have shown that mercury chloride (HgCl_2_) is very potent at producing DNA damage in mammalian cells (25,27). Despite this potency in damaging the DNA, mercury compounds have not been reported to be carcinogenic or mutagenic in many systems. Some of the mechanisms have been reviewed by which mercury induces DNA damage. New data on the effects of methyl-HgCl on DNA in cultured fibroblasts, as well as in cultured nerve cells, have also been presented (28).

UV spectroscopy describes the denaturation and renaturation of DNA in the presence of different divalent metal ions have been studied through the melting behavior of DNA (29). Hg(II) mostly prefers to bind DNA reversibly. Ultraviolet absorption spectroscopy concludes that Hg(II) binds to weekly basic nitrogene sites on the purine and pyrimidine bases. Further studies by UV spectroscopy and potentiometric methods revealed that Hg(II) interacts most strongly with AT-rich DNA. Initial Hg(II) addition is accompanied by the loss of H^+^ ions per Hg(II) ion bound (30).

High concentrations of heavy metals induce oxidative stress by increasing the formation of reactive oxygen species (ROS) like superoxide radical (O_2_), singlet oxygen (^1^O_2_) and hydrogen peroxide (H_2_O_2_) in plant cells, which are responsible for peroxidative damages to fatty acids, nucleic acids, proteins and chlorophyll (31,32). Mercury exposure causes an initial activation of T cells (33), followed by a period of T-cell proliferation in mouse models (34–36). This proliferation is more pronounced in CD4+T cells and is dependent upon the presence of adherent macrophages and the presence of IL-1 *in vitro* (at 10μM HgCl_2_) (37–39). Mercury-induced T-cell proliferation is prevented by concurrent treatment with anti-CD4 antibodies in this model (40,41). B-cell activation occurs subsequently and is dependent upon T-cell activation, although B-cell activation is directly responsible for many of the pathological outcomes of mercury-induced autoimmunity including elevated levels of circulating immune globulins like IgG and IgE. Mercury complexes have been nominated for disrupting the internal structure by breaking its DNA strands having irreparable impacts (42,43).

At the cellular level, toxicity of mercury complexes may be further explained. Mercury has been established as an extremely reactive, very soft metal, whose ions have the ability to coordinate with biologically active ligands such as carboxyl groups (40,44). This inhibition of growth may also be observed in plant cells, where the reduction in cells growth at high concentration of Hg may be correlated to high mercury accumulations by cultures. In plant cell cultures, cells have to spend extra energy to cope with the high concentration in the tissues (45). This indicates anti oxidative defense system may be involved in strategy used by these cells to survive under high mercury concentration.

In this work, mercury-complex derived from benzene-1,3,5-tricarboxylic acid has been synthesized, characterized and its cell culture study has been done for DNA damage. Through surface morphology, roughness and high compaction of surface of synthesized mercury-complex has been demonstrated. Mercury complex due to its toxic nature decides the death and life of cells focusing on cell culture and DNA damage.

## Experimental

## Materials and methods

The required chemicals were obtained from market. These chemicals were used further without purification. Mercury chloride and Benzene-1,3,5-tricarboxylic acid were obtained from Merck Chemical Company, Germany. Distilled Water, Triflouro acetic acid (TFA), Ethyl Alcohol, Methyl Alcohol, Dimethyl Formamide (DMF) and Tetrahydrofuran (THF) were used further without purification. Elemental analysis was performed on the elemental analyzer, Vario Micro Cube, Elementar Germany. FTIR spectra were recorded on Bruker Tensor 27 FT-IR spectrometer with Diamond-ATR module over the range of 4000–400 cm^−1^. XRD pattern of the mercury-complex was studied on XPert PANlytical Powder machine **Cu** Kα (l=1.54 Å) radiation. For the surface image of mercury-complex Atomic Force Microscope AFM5500M was used.

### Synthesis of mercury complex

Benzene-1,3,5-tricarboxylic acid (105 mg, 0.2 mmol) was dissolved in 5 ml of distilled water by heating on hot plate along with stirring at 200 ^0^C for 5 to 10 minutes. 1 ml of solvent tetra hydro furan (THF) was added to above solution in order to avoid the recrystallization of benzene-1,3,5-tricarboxylic acid. A solution of HgCl_2_ (135 mg, 0.2 mmol) was prepared in separate vial by dissolving in 2 ml distilled water. A clear solution was obtained upon mixing the above two solutions and sonicated for 15 minutes and then filtered. The filtrate was stand over for 10 days, and off white rod like crystals were obtained after solvent evaporation. Crystals were recovered by filtration, rinsed with mixture of water and THF *(V:V=90:10)* and dried at room temperature (Yield was 60%).

## Results and discussions

### CHNS determination

For Mercury complex (C_18_H_10_HgO_12_) (M.W=618.55amu)

|  |  |  |  |
| --- | --- | --- | --- |
| Anal.Calcd., % | C, 31.53 | H, 3.82 | N, 6.02 |
| Found, % | C, 31.29 | H, 3.46 | N, 1.61 |

The ligand (Benzene-1,3,5-tricarboxylic acid) has percentage values determined theoretically (C, 46.2%, H, 17.00%). The percentage values of C,H in ligand have been decreased in mercury complex, that indicates the binding of ligand with metal ion.

### FTIR of mercury complex

The arrangement of peaks in mercury-complex are as; 740 cm^-1^ (sharp), 1255.38 cm^-1^ (broad), 1322 cm^-1^ (sharp), 1539cm^-1^ (broad), 1637 cm^-1^ (sharp), 2847.11 cm^-1^ (medium and broad). Broad peak at 2847 cm^-1^ indicates the presence of (–COOH) group and peak at 1637 cm^-1^ shows the binding of metal with ligand (TMA). These lower values of C=O indicates that ligand is coordinated in boning of mercury complex. The specific peak at 1322 cm^-1^ must be due to stretching vibration of C=O bond of carboxyl group on aromatic ring. While peaks at 3688 cm^-1^ and 3578 cm^-1^ represent –OH group from ethanol.

The theoretical structure of mercury-complex has been in a square planar geometry, mercury is coordinated by two oxygen atoms [O(14) and O(15)] of ligand (TMA) and two oxygen atoms [O(23), O(24)] of another ligand (TMA) in [Hg(TMA)_2_]. The molecular mass of proposed complex is 618.55 amu. Empirical formula of complex is C_18_H_10_HgO_12._

### Microscopic imaging

The morphology of the synthesized mercury-complex has been analyzed by microscope. Images were recorded in the range of 2-5 mm with magnification of 200-500 x. Microscopy Imaging is the technique where a focused beam of light is used and gives us the information about microscopic morphology. The surface/shape information of synthesized complex has been observed under microscopic analysis. The sample was fixed in the specimen stub by covering it with glass slide cover. Then the microscope was set to get the high-resolution image of the complex. In images, we can see the homogenous dispersion of rod like crystals; these crystals are not strongly agglomerated because of their low surface energy. Due to low aggregating or overlapping of smaller particles there are a few larger crystals. The microscopic pictures clearly show randomly distributed crystals with smaller size and it is observed that the homogeneous rod-shaped crystals and flakes like structure. Following are the results of our microscopic analysis.

### Atomic force microscopy

Atomic-force microscopy (AFM) is a powerful technique that can image almost any type of surface, including polymers, ceramics, composites, glass, and biological samples (46). AFM is used to measure and localize many forces, including <u>adhesion strength,</u> magnetic forces, and mechanical properties. AFM is performed using a sharp tip about 10–20 nm in diameter attached to a cantilever. To study mercury-complex, the most versatile and powerful microscopic technique is certainly atomic force microscopy. It is adaptable and multipurpose because AFM can not only show 3D topographic images but also provides detailed surface dimensions with calculations and measurements (47). AFM is considered a powerful tool because images produced are at atomic level in angstrom scale, providing not only height details but only small amount of sample is required. To scan a sample for topography, a cantilever is used having a highly sharp tip. As the tip of cantilever approach the complex material, the tip is attracted towards the sample and this attraction deflects the tip to sample. Though, when the cantilever tip touches the sample, a repulsive force is generated that deflects the tip away from the sample surface. The deflection of cantilever is recorded through a laser beam on deflection toward or away from sample surface. The slight changes in cantilever i.e. deflection is detected through a reflected beam (48).

Surface morphology of synthesized mercury-complex was examined through AFM5500M. Fig 5 shows the surface image of mercury-complex. The phase topography of the selected section as measured is displayed as a surface plot, with the calculated depth of the marked pattern scale in absolute units. The size of the measured area is 5 × 5μm^2^. Here the phase profile is plotted as a surface plot with absolute depth values in μm as obtained by the numerical processing. Obviously, the contrast of the height of the inner parts of the sample decreases if the spatial resolution comes to its limit, which is the case for structure sizes below 1 μm. However, for larger structures, the contrast and the measured phase profile are independent from the shape or the location of the structure. It shows the roughness of synthesized material. These images showed the high compactness of material synthesized.

**Fig 1.**
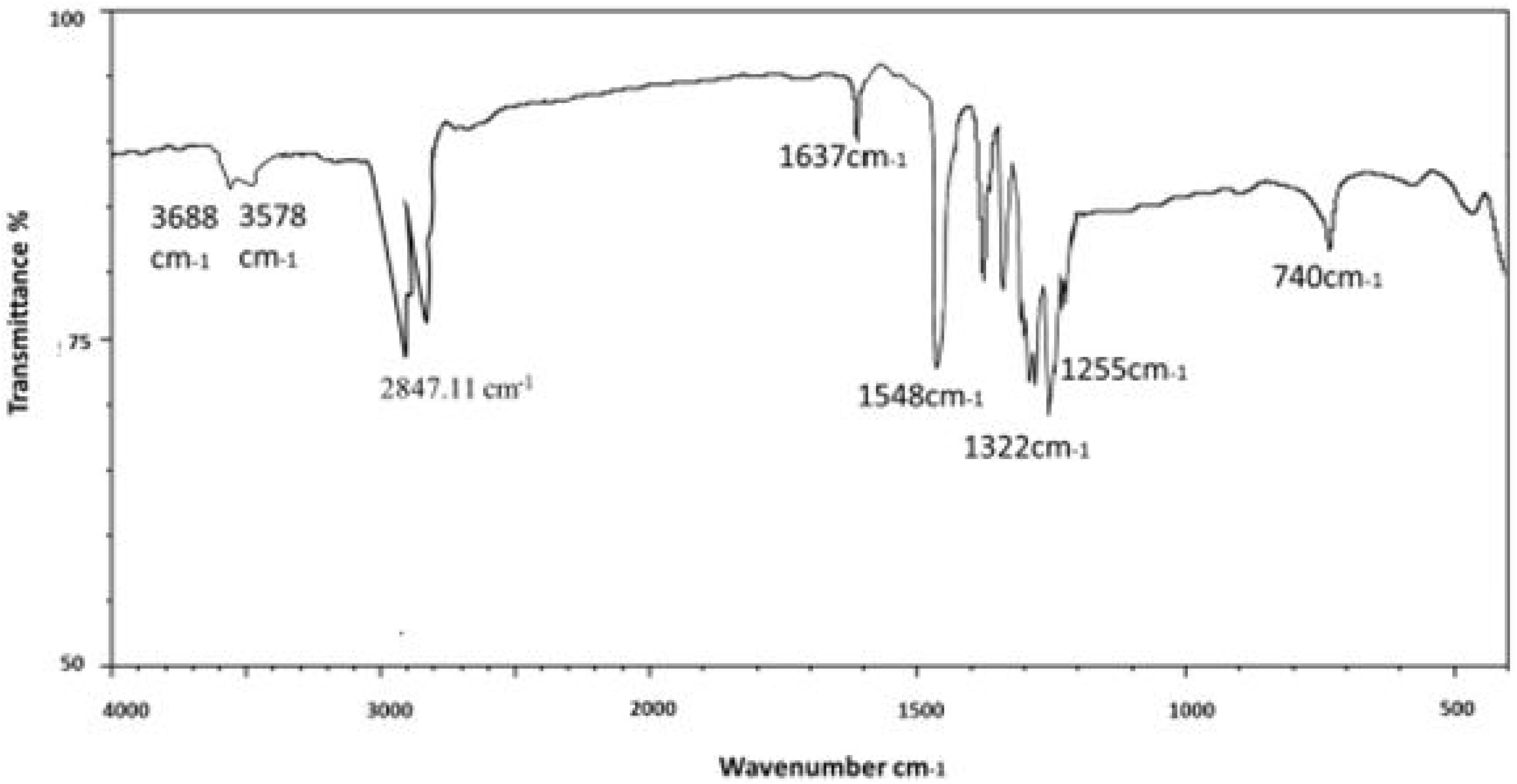
FTIR spectra of mercury complex.

**Fig 2.**
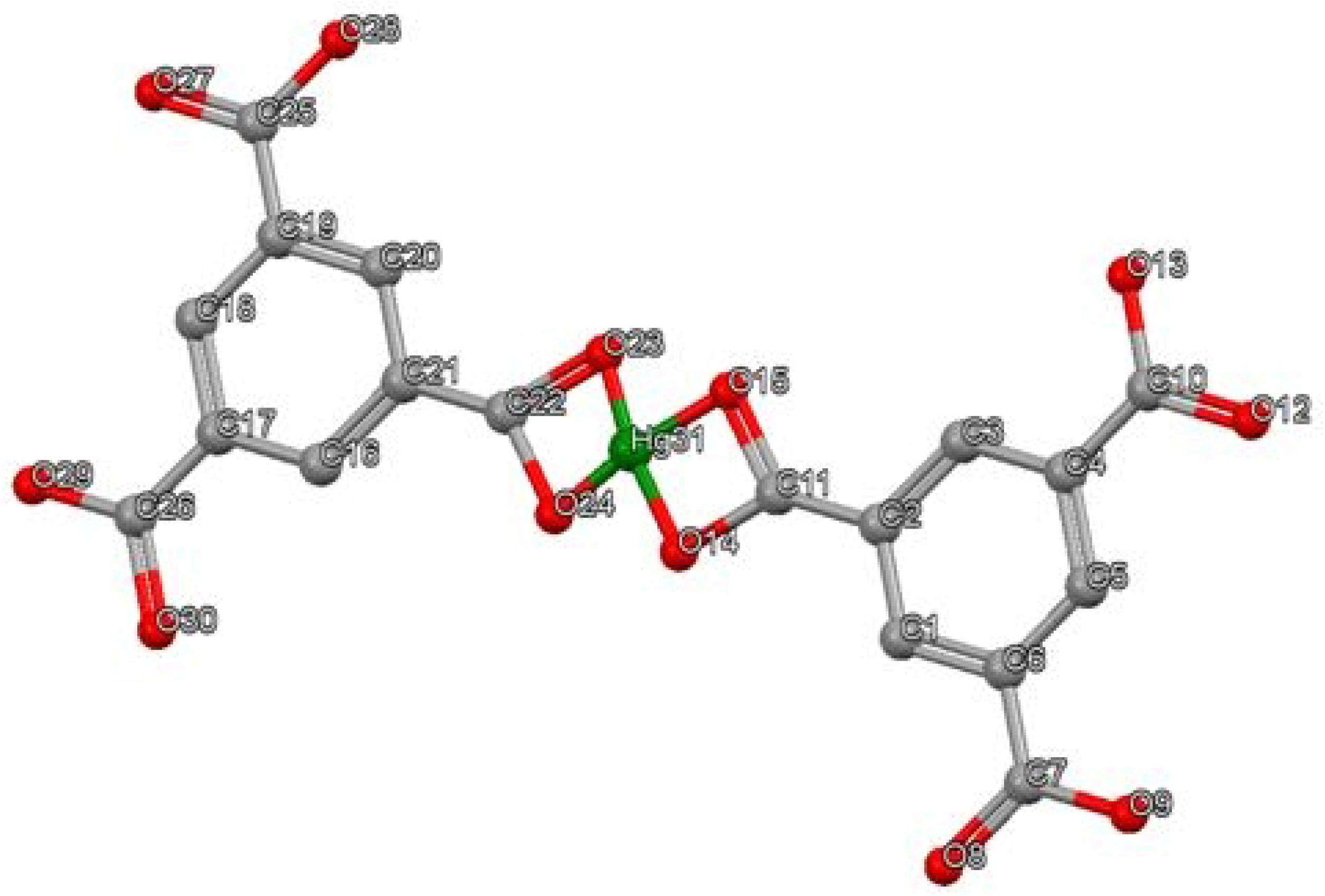
Proposed structure of mercury complex [Hg (TMA)_2_].

**Fig 3.**
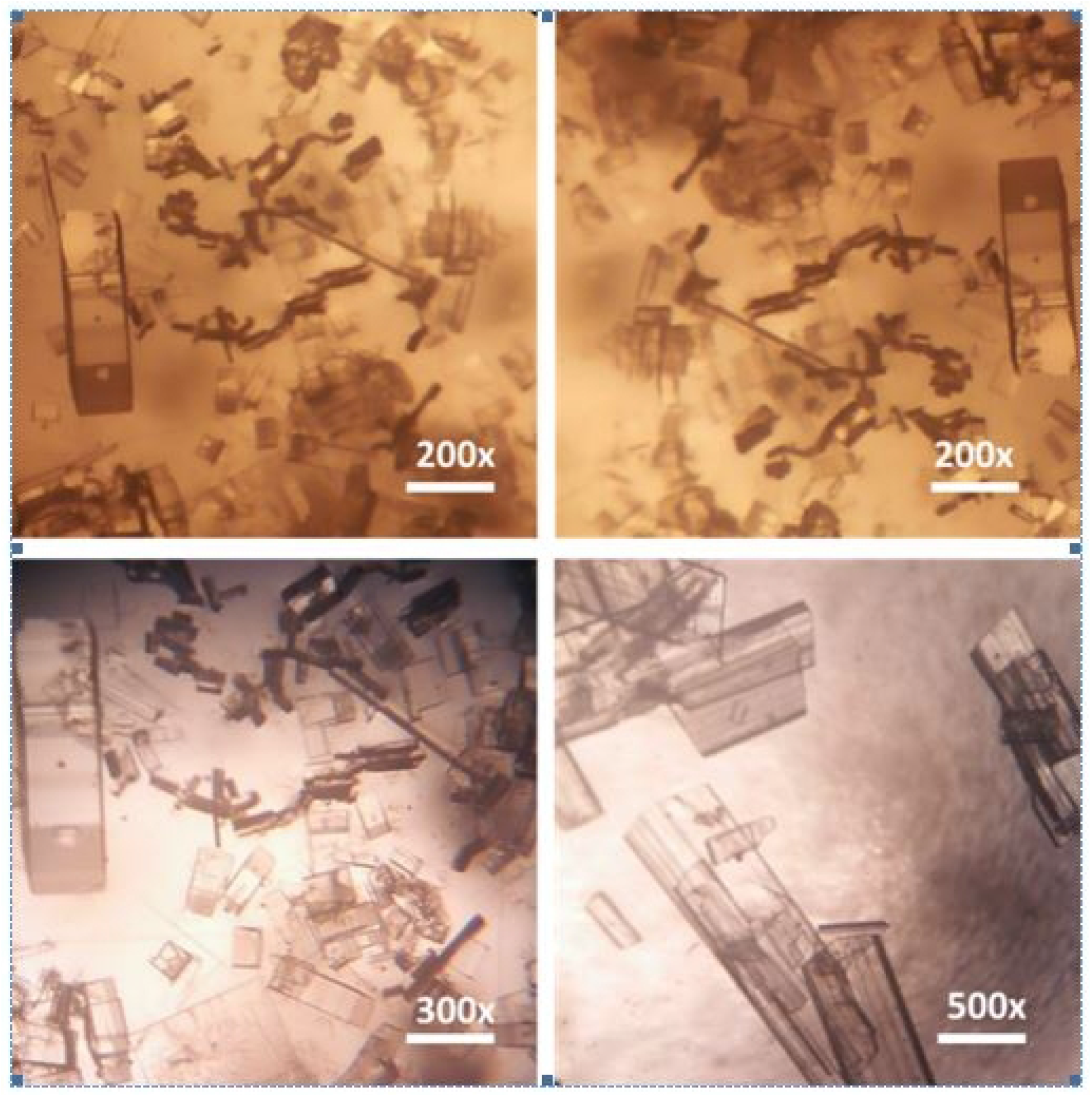
Microscopic imaging scans of Hg-complex in the range of 2-5 mm with magnification of 200-500x.

**Fig 4.**
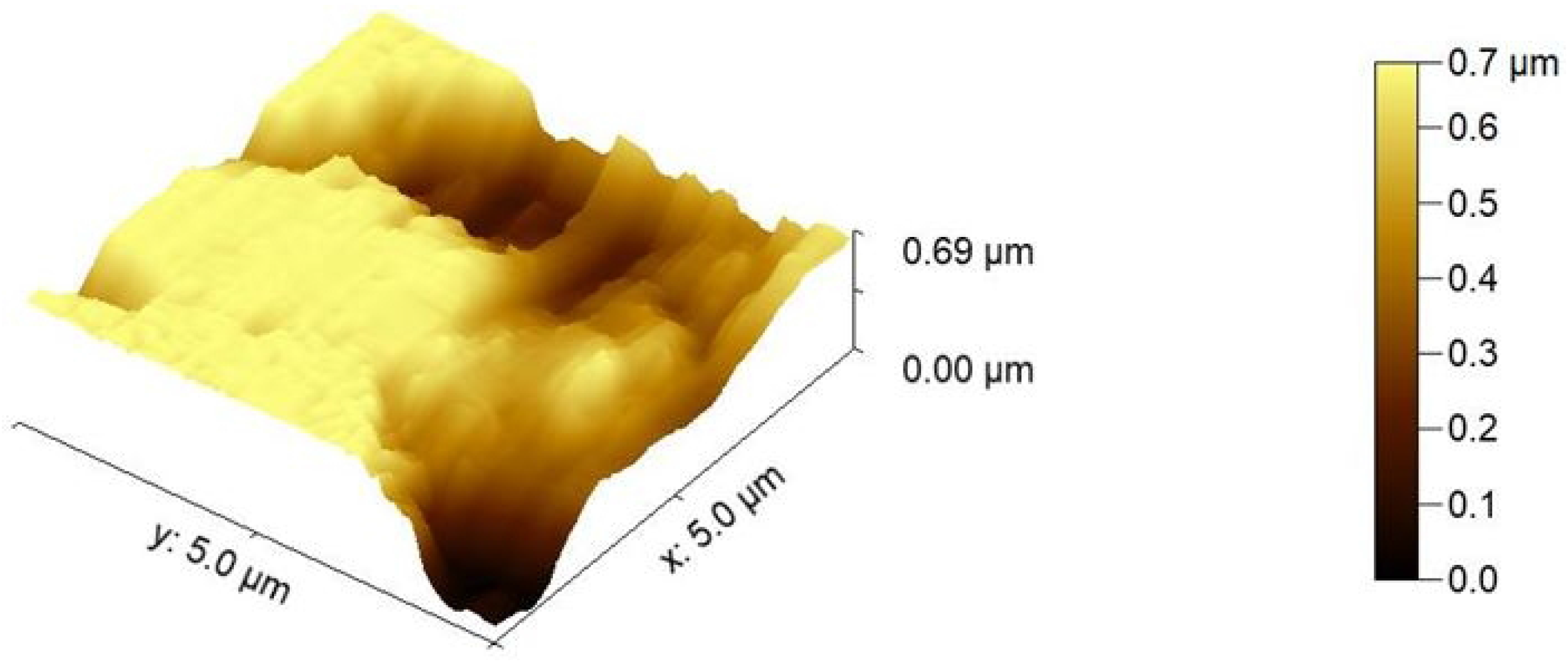
Atomic force microscopy scans of Hg-complex. The phase topography of selected section as measured with the phase method is displayed as a surface plot of the phase profile, displaying the absolute calculated phase shift in microns.

**Fig 5.**
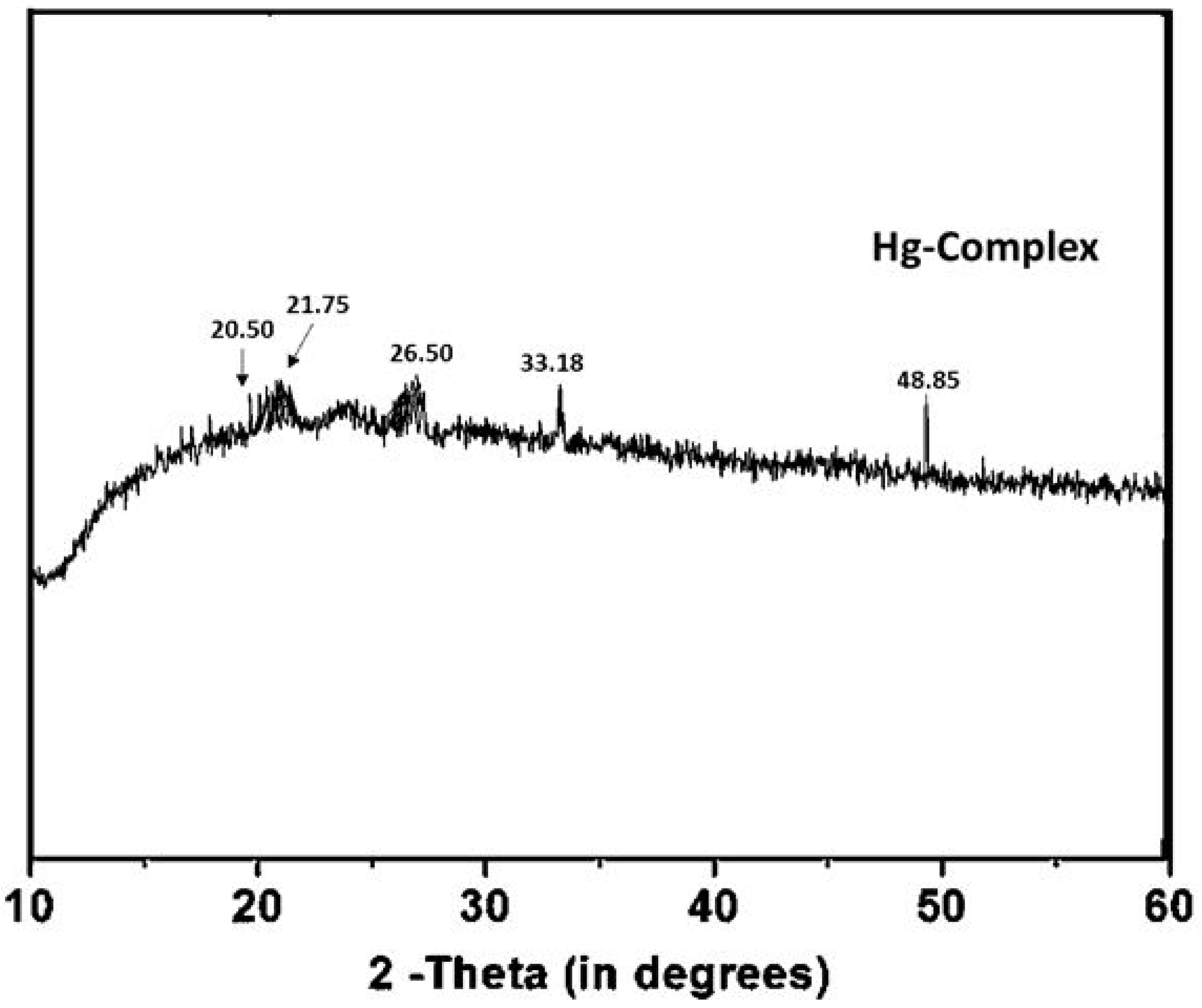
XRD analysis of mercury complex.

Atomic force microscopy (AFM) of DNA has changed since the early images of Albrecht Weisenhorn (46). In the hope of sequencing DNA with the AFM, we started by imaging single-stranded DNAs only 20 bases long, which were not only very small but also very easily pushed by the AFM tip with the methods of sample preparation and imaging available at that time (49). Two advances created a major breakthrough: 1) imaging plasmid DNAs, which have characteristic circular shapes of known lengths, and 2) imaging in dry air, which helps to keep the DNA well bound to the substrate (50,51).

### X-Ray diffraction (XRD) analysis

The XRD pattern of the mercury-complex was studied by XPert PANlytical Powder machine **Cu** Kα (l=1.54 Å) radiation. The crystalline sample was well dried and grinded to make fine particles prior to XRD analysis (52). The XRD patterns of mercury-complex confirming the as synthesized complex is pure in phase, having no other phase with no detectable impurities. The XRD spectrum of the mercury-complex is shown in Fig 5. The sample had diffraction peaks at 2θ = 20.5°, 21.75°, 26.5°, 33.18° and 48.85 which were attributed to literature peaks based on face-centered cubic structures of mercury-complex, when compared to library references (53,54). It affirmed the composition of the complex containing mercury.

### Cell culture study

To study the effect of mercury-complex on cell culture, complex was investigated for effects. For this purpose, cells were cultured in the absence and presence of mercury-complex, with varying concentrations after three days. The branching and over lapping of dead cells and live cells were counted in fluorescence imaging. The effects of mercury-complex with different concentrations were studied. The increase in concentration of mercury-complex augmented the effects of mercury over cell cultures. The cell death, over lapping of cells and margins diminishing, is observed in case of mercury-complex presence compared to absence of complex. “A” image represents control i.e. without mercury-complex while “B” represents cell culture with presence of mercury-complex (Fig.6A&B). The arrow represents the major area of cell culture effected/damaged by mercury-complex. The Fig 7 shows the results of cell culture study after 3 days exposure of mercury-complex, which indicates a lot of collateral damage and death of cells.

**Fig 6.**
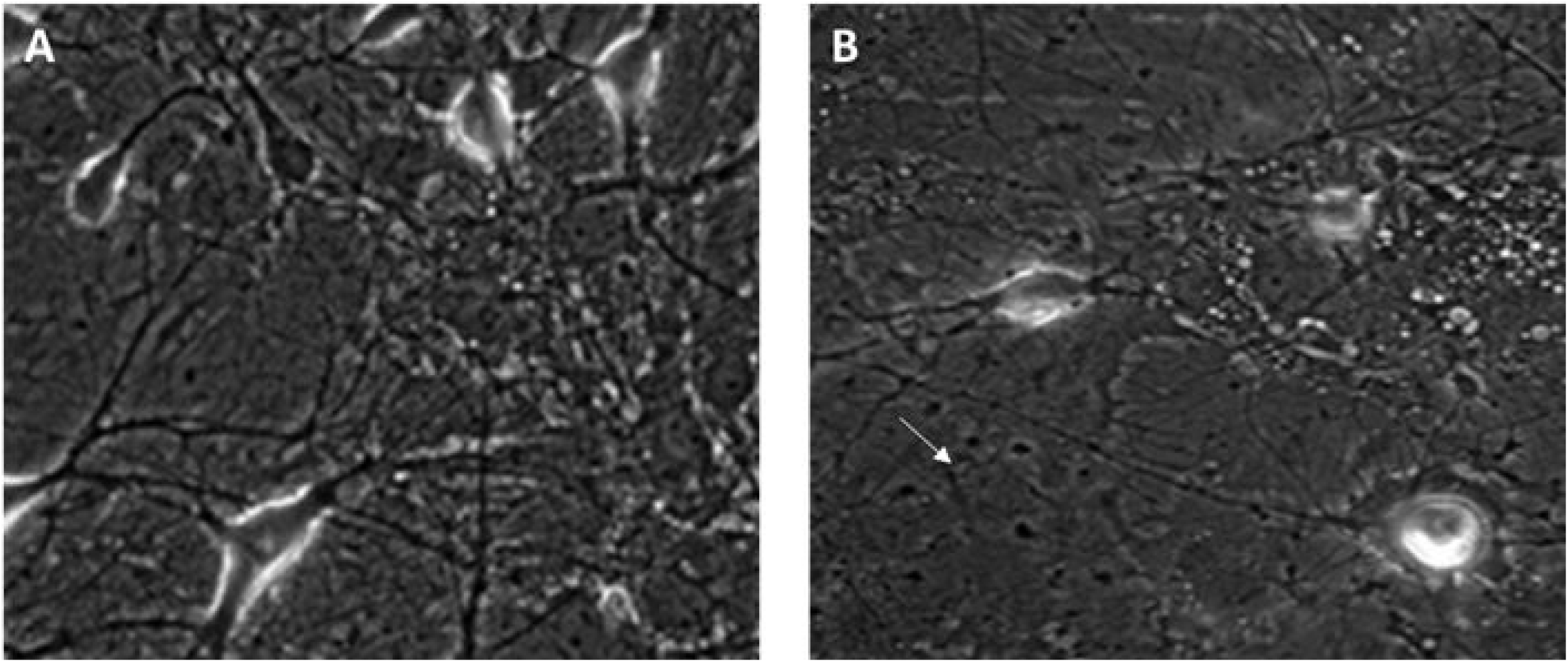
Cell culture study; A) Control cell culture without Hg-Complex and B) Cell culture in presence of Hg-Complex.

**Fig 7.**
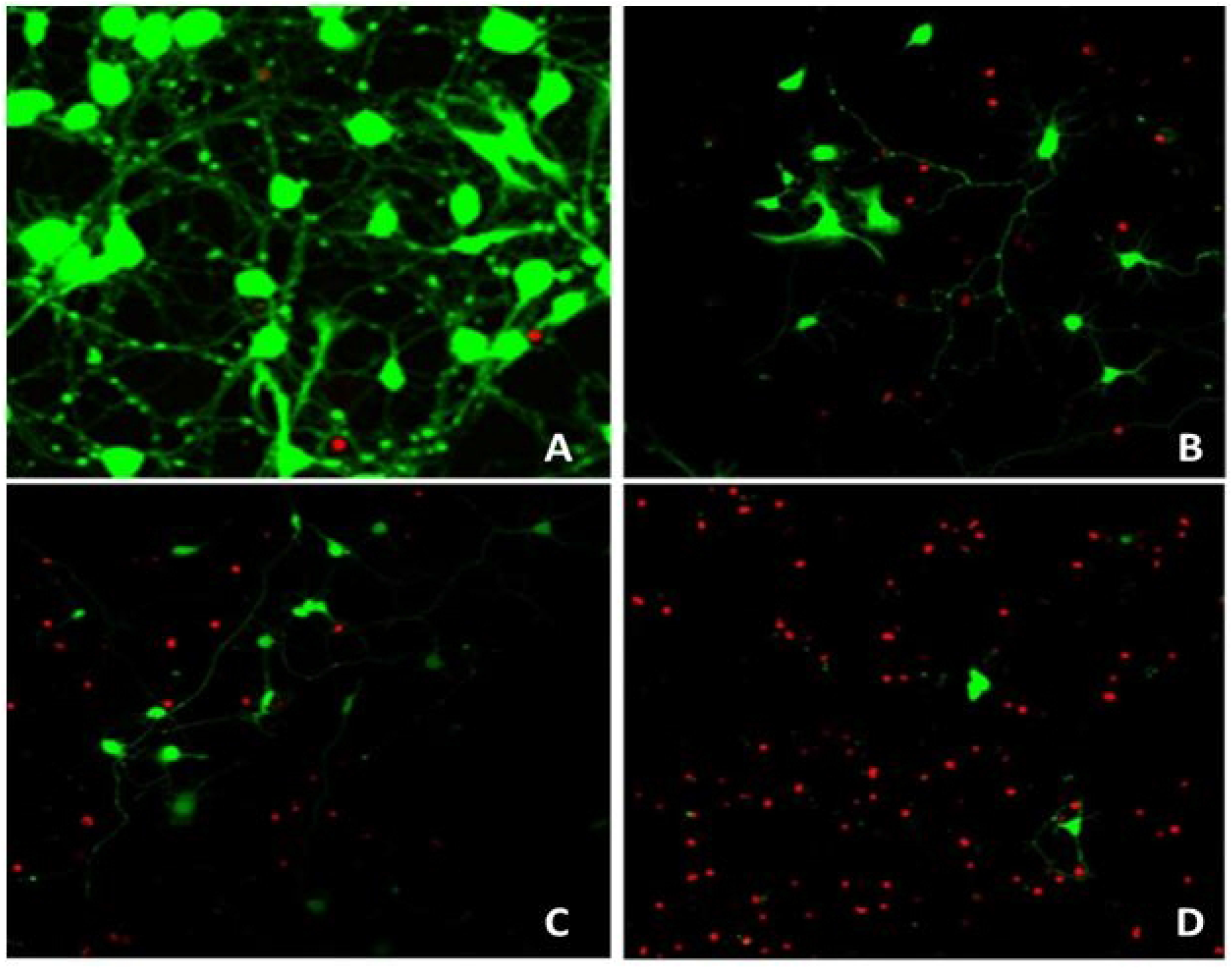
Live/Dead cell assay; A) Control without Hg-complex and B) Cell culture study 1 μg/ml, C) 10 μg/ml and D) 50 μg/ml; in presence of Hg-complex.

Despite the clustering increase compared to control, a number of cell populations die in presence of complex. With increase in concentration of mercury-complex the count of cell death increases and living cells number fall drastically (Fig7).

### Live/Dead cell assay

To further quantify the effects of mercury complex on cell viability, we subsequently performed live/dead assay through fluorescence imaging in presence or absence of mercury-complex for 3 days using concentration of 1μg/ml, 10μg/ml and 50μg/ml. Fig 8A represents live or dead cell assay without mercury complex, whereas Fig 8(B,C,D) explains cell assay with different concentrations of mercury complex. Fig 9 is the graphical elaboration of cell assay at different concentrations.

**Fig 8.**
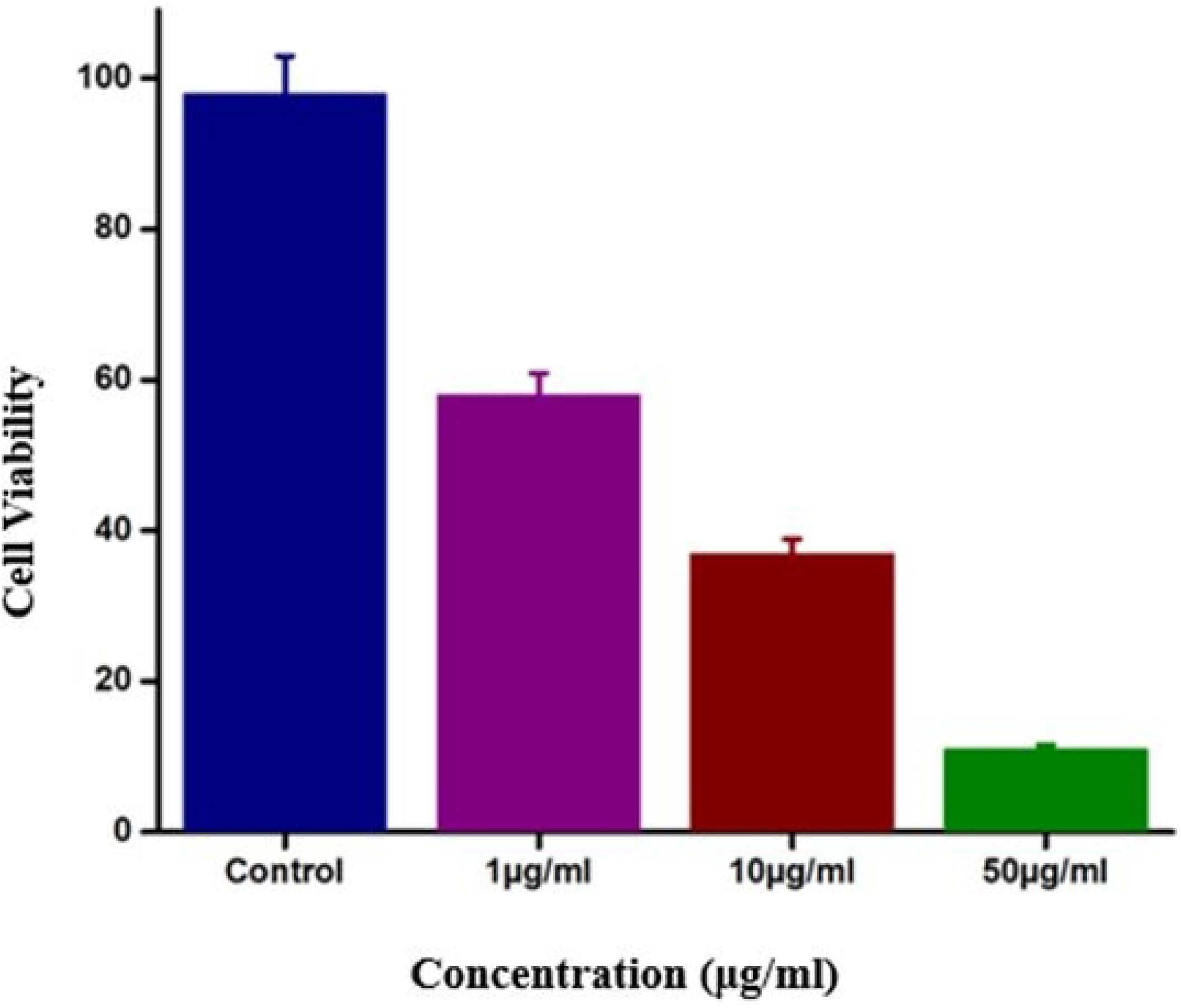
Live/Dead cell assay graph; Control without Hg-complex and 1 μg/ml, 10 μg/ml and 50 μg/ml; in presence of Hg-Complex.

Mercury complexes have damaging effects on cell growth and culture; these may be used for further medical treatment of different diseases like cancer. It is assured, that mercury complexes have the ability to stop the growth of cells. The images of cell growth and death could be verified by using a fluorescence microscope. Fluorescence signals may be localized in the intracellular area, indicating a sub cellular distribution of mercury complex (8,55).

Presence of Hg complex damages the cell as shown in the research work by Kim and Yuan (56,57) that can be compared to the mercuric or methyl-HgCl_2_ ion, because of its high reactivity and is capable of producing very striking damage to cells at low concentrations. It has also been justified that the mercury ion has the capacity to generate oxygen radicals in cells. The mechanism has not been uncovered that how this occurs, but it has previously shown that it occurs at 40C^0^ to a somewhat lesser extent than at 37C^0^(43,58). Additionally, HgC1_2_ treatment of intact cells has been shown to induce strand breaks in nucleoids, isolated from cells at the same concentration as it induces strand breaks when added directly to the nucleoids *in vitro*. The present research results are aligned with study conducted by (28) that with the increase of concentration and time more damage of cells occur with the ability to target the focused area of cell culture (59,60). Its high accumulation in the human body causes prenatal brain damage, serious cognitive and motive disorders, vision and hearing loss, and even death. So there should be the best analysis that mercury-complex may be utilized for medical treatment keeping in mind its hazardous nature as well (8,61).

Under controlled conditions with high accuracy the mercury-complex has the ability to damage the affected cells by controlling its growth with change its concentration and time. According to our knowledge, this is novel mercury-complex based on atomic force microscopy explanation, has the ability to damage cells. It could be further utilized for treatment of cancer affected cells and other diseases.

The development of multifunctional micelles with unique chemical and physical properties conferred by pH-sensitive uracil-mercury-uracil (U-Hg-U) linkages and tunable structural and dynamical features due to the presence of hydrogen-bonded uracil moieties represents a significant breakthrough in the construction of mercury-containing supramolecular polymers. Importantly, in vitro experiments showed that incorporating the U-Hg-U complexes into the micelles not only increased the efficiency of selective uptake via endocytosis into cancer cells, but also accelerated the induction of massive apoptotic cell death (62). As a result, this research provides critical new insight for the development of metallo-supramolecular polymeric micelles that may significantly improve the safety and efficacy of cancer therapy and bioimaging without the use of anticancer drugs or fluorescent probes.

Indeed, metal-polymer coordination-based nanoparticles could be used to create multifunctional nanoplatforms that enable tumor-selective delivery of therapeutic agents to suppress tumor cell proliferation, or they could be used as diagnostic tools for early tumor detection or assessing the pharmacokinetics, pharmacodynamics, and efficacy of anti-cancer drugs (63).

## Conclusion

Today, a lot of research has been progressed for the use of bio-compatible materials, the synthesis of complexes which includes the use of organic components or the compounds obtained from living organisms. The main goal of this research was to investigate the effects of mercury-complex derived from benzene-1,3,5-tricarboxylic acid on cell culture. For this purpose, we synthesized mercury-complex via sonication, characterized and measured its potential effects on cell culture. Mercury based complexes are suggested not to be used as eco-friendly and economical agents, but this mercury complex is proved effective through cell culture study. At low concentration, its damaging effects are low but when we increased its concentration, it completely damaged the cell culture as confirmed by cell viability. For all the mercury complexes under study, DNA damage can be attributed by the strand breaks of mercury complexes. Mercury-complex is non-biocompatible, non-biodegradable, and toxic complex, which has demonstrated damaging effects against cell growth. The growth activities of mercury-complex is dependent on numerous factors such as type of cells used in study for cell culture, molecular weight (MW) and concentration of mercury-complex provided. The mercury-complex has showed its damaging effects on cells with increase of concentration and time. The mercury-complex is biologically more active and efficient for medical treatment.

## Acknowledgements

Authors are thankful to Materials Chemistry Laboratory, Department of Chemistry Government College University Lahore for providing research facilities.

## Author Contributions

**Javed Hussain Shah**: Conceptualization, Methodology, Software, Investigation, Visualization, Writing-Original draft. **Shahzad Sharif:** Conceptualization, Validation, Writing-Review and Editing, Investigation, Supervision, Project administration. **Rashid Rehman:** Data curation, Formal Analysis. **Anum Arooj:** Software, Validation, Writing-Review and Editing.

